# Genetic surveillance of target site mutations and copy number variants associated with the escalating insecticide resistance in *Anopheles gambiae* (s.l.) in Ghana

**DOI:** 10.64898/2026.10.14.759058

**Authors:** Anisa Abdulai, Isaac Kwame Sraku, Christopher Mfum Owusu-Asenso, Abena Ahema Ebuako, Yaw Akuamoah-Boateng, Abdul Rahim Mohammed Sabtiu, Jemima Boateng, Christabel Koranteng, Bright Churchill Obeng, Godfred Amoateng, Fred Aboagye-Antwi, Patrick F. Ayeh-Kumi, Yaw Asare Afrane

## Abstract

The spread of insecticide resistance in malaria vectors poses a major challenge to control programs. As insecticide-based interventions intensify, selection for resistance alleles increases, making it essential to understand the genetic mechanisms underlying resistance in local *Anopheles* populations. This study characterized target-site resistance mutations and copy number variants (CNVs) in *Anopheles gambiae* s.l. from nine sites with different malaria vector interventions and ecological contexts. WHO bioassays revealed moderate-to-high pyrethroid resistance (4–65%) at all sites, while Obuasi (IRS) showed markedly elevated organophosphate resistance (1.7%) compared to other locations. The L995F *kdr* mutation was strongly associated with pyrethroid resistance (*P* < 0.05) and occurred at high frequencies in IRS and standard LLIN sites (0.82–1.0). Additional *kdr* variants (L995S, N1570Y, I1527T, V402L) appeared at variable frequencies (0–0.59). The *Ace-1* G280S mutation was significantly associated with pirimiphos-methyl resistance (OR=2.20, P<0.001). *Ace-1* gene duplications occurred in 22.8% of mosquitoes, predominantly in individuals carrying the G280S resistance allele (P<0.001). Carboxylesterase CNVs were also identified, with *Coeaxf*_Dup1 (30.4%) significantly associated with pirimiphos-methyl resistance (OR = 1.68, *P* = 0.02). These findings demonstrate the widespread, multifaceted resistance mechanisms in *An. gambiae* s.l. across Ghana and highlight the need for locally tailored vector control strategies.

## Introduction

Malaria is a major public health issue in sub-Saharan Africa (SSA) (1). Vector control has been a crucial part of all malaria control and elimination efforts (2). The current first-line vector control methods rely on long-lasting insecticidal nets (LLINs) and indoor residual spraying (IRS)(3). These tools played a major role in the significant reduction in malaria transmission mainly in sub-Saharan Africa over the last two decades (4). Pyrethroids are the most commonly used insecticides and are currently used in all LLINs (5,6). Insecticides like organophosphates and carbamates are mostly used for IRS (7). However, the widespread use of insecticide-based vector control tools in SSA have led to a selection pressure for the development of insecticide resistance in malaria vectors through natural selection (8–10).

Knockdown resistance (*kdr*) mutations in the target site for pyrethroids, that is, the voltage-gated sodium channel (*vgsc*) such as L995F (previously known as L1014F) and L995S (previously known as L1014S|), occurring on the same codon, have been associated with pyrethroid resistance in *Anopheles gambiae* s.l mosquitoes (11–14). Another *kdr* mutation, N1570Y significantly increases pyrethroid resistance in African malaria vectors when in combination with the L995F mutation (15). Other non-synonymous mutations have been found to occur almost exclusively on the haplotypes carrying the known L995F *kdr* mutation and may intensify or compensate for the L995F resistance genotype (16). Notably, a novel mutation, I1527T was found in strong linkage with another novel mutation, V402L and has been associated with pyrethroid resistance in *An. gambiae* mosquitoes and other insect species (16,17).

Historically, organophosphates (OPs) resistance has been mainly linked to the *Ace-1* locus, where a complex interplay of either heterogeneous or homogeneous gene duplications, along with the non-synonymous *Ace-1* G280S mutation, contributes to varying levels of resistance in *Anopheles gambiae* s.l mosquitoes(18–21). Studies have shown that increased copies of *Ace-1* and the presence of the *Ace-1* G280S mutation is driving pirimiphos methyl resistance in West African *Anopheles* populations(22,23). Currently, there is high selection pressure for organophosphate resistance in vectors because of the increase in the use of OPs for malaria vector control and in agriculture (24,25). The occurrence of these genetic variants may be driving increasing OPs resistance in Anopheles mosquitoes in Ghana and more research is needed to understand the genetic mechanisms of organophosphate resistance in *Anopheles gambiae* mosquitoes in Ghana.

Overexpression of major detoxification enzymes such as cytochrome p450s (CYP450s), glutathione-s-transferases and esterases also mediate insecticide resistance in malaria vectors (9). Cytochrome p450s overexpression especially CYP6 and CYP9 families has been mainly linked to pyrethroid resistance in *Anopheles gambiae* mosquitoes (26,27). Overexpression of esterase genes have been majorly linked to organophosphate and carbamate resistance (28,29). Copy Number Variations (CNVs) of detoxification genes have been associated with the increase in metabolic resistance in Anopheles mosquitoes. Duplicated Cytochrome p450s such as *CYP6P9a/b* and *CYP6AA1* genes to be significantly associated with pyrethroid resistance (17,30). Another study by Nagi *et al*. (21) recently found duplicated esterase gene (*Coeae1f* and *Coeae2f*) to be driving resistance to pirimiphos-methyl in West African *An. gambiae*.

The increase in the frequencies of genomic variants associated with insecticide resistance may significantly impact the success of current insecticide-based control tools such as LLINs and IRS. Information on the distribution of insecticide resistance mutations and CNVs in *Anopheles gambiae* populations in Ghana is limited. There is a need for more genetic surveillance of insecticide resistance alleles to help in the development of effective insecticide management. Here, we investigated the distribution of target site resistance mutations and CNVs in *Anopheles gambiae* sensu lato (s.l) mosquitoes from sites with different vector control interventions in Ghana.

## Methods

### Study Site

The study was conducted in 9 sites categorized according to the vector control interventions currently being employed in those sites. These sites include indoor residual spraying sites (Kpalsogu and Obuasi), Pyrethroid-PBO nets sites (Tolon), chlorfenapyr nets (Takoradi and Elubo) and standardized LLINs sites (Libga, Konongo, Dodowa and Ada) (Fig 1). These sites are part of the Ghana National Malaria Elimination Programme (NMEP) sentinel sites and spread across the three main ecological zones of Ghana. Ada (5° 46’51.96 "N 0° 35’12.84 "E),

**Fig 1:**
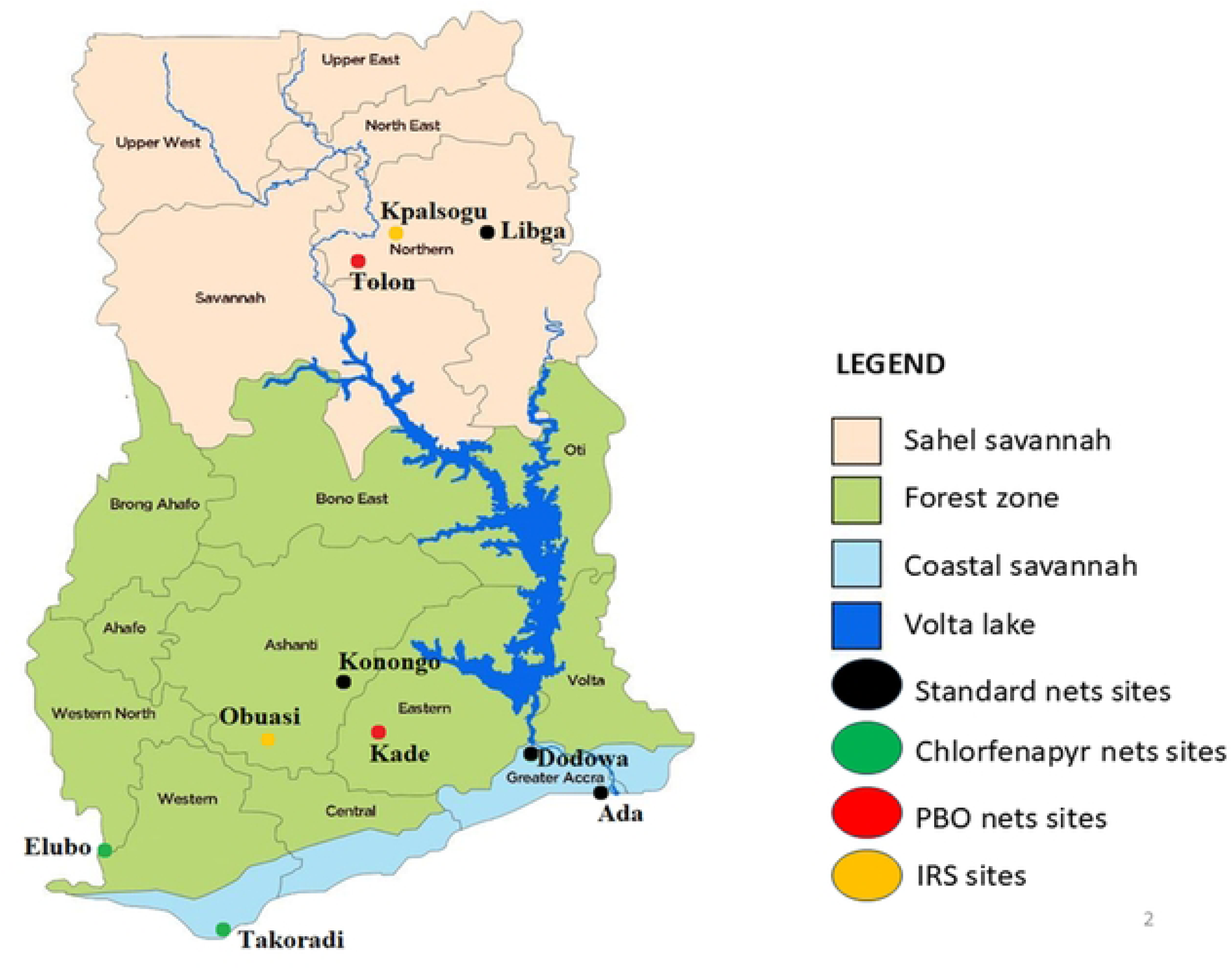
Map of the sites used for this study.

Takoradi (4.9016° N, 1.7831° W) and Elubo (5° 16’ 60.00" N, 2° 45’ 59.99" W) is located in the coastal savannah zone of Ghana. Irrigated farming activities occurs in Ada and have created many suitable breeding sites, thus increased the densities of mosquitoes in the area (31). Takoradi is a port city in the south-western part of Ghana. Elubo is a town located near the border with Cote d’Ivoire. Konongo (6° 32’3.05 "N 1° 14’42.22 "W) and Obuasi (6.2012° N, 1.6913° W) are located in the forest zone of Ghana. These areas have a wet semi-equatorial climate characterized by two distinct rainy seasons: a long rainy season from May to July and a short rainy season from September to November. Indoor residual spraying is carried out in Obuasi. Kpalsogu (9° 22’33.34 "N 0° 42’29.67 "W), Tolon (9.4309° N, 1.0646° W), and Libga (9°35′32.26"N, 0°50′48.8"W) are districts located in the Savel Savannah zone of Ghana. They have a unimodal rainfall pattern from May to November and a long dry season from December to April. The mean annual temperature is around 28°C but can get to a maximum of 42°C. Irrigated farming is practiced in these rural communities throughout the year, creating numerous breeding habitats for *Anopheles* mosquitoes. Indoor residual spraying and seasonal malaria chemoprevention (SMC) are carried out in Kpalsogu.

### Larval Mosquito Sampling

Anopheles larval samples were collected from breeding sites across the different study sites using WHO 350ml standard dippers. Anopheles larvae were transported to the insectary at the Department of Medical Microbiology, University of Ghana Medical School, Accra, for samples collected in the coastal zone, where they were raised to adults. Anopheles samples collected in the forest and sahel zone were transported to the insectaries of AngloGold Ashanti malaria control programme (AGAmal) in Obuasi and President’s Malaria Initiative (PMI) Project Office in Tamale respectively. The larvae were fed on TetraMin^®^ fish meal and maintained at 27 ± 2 °C. Emerged adults were fed on 10% sugar solution. The F0 adult mosquitoes were used for WHO insecticide resistance testing and subsequent analysis.

### WHO Susceptibility tests

Four replicates of 20 nonblood-fed female *Anopheles gambiae* s.l mosquitoes (aged 3 to 5 days) were exposed to papers impregnated with pyrethroids {type I permethrin (0.75%) and type II deltamethrin (0.05%)} and organophosphate {pirimiphos-methyl (0.25%)} for 1 hour according to WHO criteria to determine phenotypic resistance(31). Two replicates of 20 female mosquitoes were also exposed to oil-impregnated papers as controls. Mosquitoes in both the test tubes and the control tubes will be exposed to the insecticide-impregnated papers for 1 hour. Knockdown was recorded at 10-minute intervals over a 1-hour exposure period. After exposure, mosquitoes were transferred to holding tubes and maintained on 10% sucrose solution. Mortality was assessed after a 24-hour recovery period, and resistance status was determined according to WHO guidelines(31). The Kisumu strain of *An. gambiae* was also used as a susceptible control to verify the efficacy of the insecticide-treated papers. Both dead (susceptible) and surviving (resistant) mosquitoes were preserved in eppendorf tube containing silica gel for subsequent molecular analysis.

### Morphological and Molecular Species Identification

All mosquitoes collected were morphologically identified using morphological keys by Gilles and Coetzee(32). Members of the *An. gambiae* s.l. complex was further identified by polymerase chain reaction (PCR) using well established protocols by Scott *et al*.(33). Restriction Fragment Length Polymorphisms (RFLP) was used to distinguish between the sibling species, *An. gambiae s.s* and *An. coluzzii* using protocols from Santolamazza *et al*.(34).

### Genotyping of Knockdown Resistance Mutations in *An. gambiae* s.l mosquitoes

The presence of knockdown resistance (*kdr*) (L995F, L995S, N1570Y, V402L and I1527T) mutations were detected in pyrethroid resistant and susceptible *An. gambiae* s.l mosquitoes using conventional and real-time PCR assays. The *kdr* mutations L995F and L995S were detected using allele-specific PCR using protocols by Bass *et al*.(35). The kdr mutation N1570Y mutation was detected using TaqMan probe-based assays by Jones *et al*.(15). Locked nucleic acid (LNA) based real-time PCR assays were to detect two novel mutations V402L and I1527T using established protocols by Williams *et al*.(36).

### Detections of Ace-1 G280S mutation and Ace-1 gene duplication in An. gambiae s.l mosquitoes

*Anopheles gambiae* s.l mosquitoes exposed to the Pirimiphos methyl were genotyped for *Ace-1* G280S mutation (38). The presence of duplications in the *Ace-1* locus was detected in *An. gambiae* s.l mosquitoes that were resistant or susceptible to the Pirimiphos-methyl using PCR-based protocol by Assogba *et al*.(37). A pair of forward and reverse primers Agduplispedir2 and AgduplispeRev1) were used to amplify the overlapping region of the duplicated junction containing the *Ace-1* locus (*Ace1*_AgDup) using PCR. The resulting 460 bp fragment spans the junction and is thus amplified only in individuals with multiple copies of the *Ace-1* locus.

### Detections of Carboxylesterase gene duplications in *An. gambiae* s.l

Two gene duplications containing the two alpha-esterases, *Coeae1f* and *Coeae2f* locus, which are associated with pirimiphos-methyl resistance were also detected by PCR using well-described protocols by Nagi et al.(21). A set of three primers (Forward, reverse and reverse control primers) each were used to detect duplications junctions (for Coeaexf_Dup1_2 and Coeaexf_Dup2_2) spanning the *Coeae1f* and *Coeae2f* locus PCR assays. Products were visualized on 2% agarose gels. Bands of 178 bp and 167 bp was indicative of a positive sample for Coeaexf_Dup1 and Coeaexf_Dup2 respectively. Band sizes 429bp and 400bp was indicative of control bands for Coeaexf_Dup1 and Coeaexf_Dup2 respectively.

### Data Analysis

Data was represented as tables and figures. Insecticide susceptibility tests were interpreted according to WHO guidelines(31). Mosquito populations with mortality rates of 98% to 100% were classified as susceptible, while mortality rates of 90% to 97% were classified as suspected resistance and mortality rates below 90% were classified as resistant. Chi-square tests or Fishers’ exact tests were used to assess significant associations in phenotypic resistance and the presence of resistant mutations across study sites, interventions and phenotypes. Odds ratios were calculated to determine probabilities and p-values less than 0.05 were represented as statistically significant.

## Results

### Species discrimination of *Anopheles gambiae* s.l across the study sites

Out of a total of 380 *An. gambia*e s.l mosquitoes that were molecularly identified, *Anopheles coluzzii* was the most predominant species (301, 79.2%), followed by *Anopheles gambiae* s.s (60, 15.8%), *Anopheles arabiensis* (10, 2.6%) and *An. gambiae* s.s/*An. coluzzii* hybrids (9, 2.4%). *Anopheles coluzzii* was the common species in all the study sites except in Tolon, where *An. gambiae* s.s was the predominant species (47.5%) (Table 1).

**Table 1:**
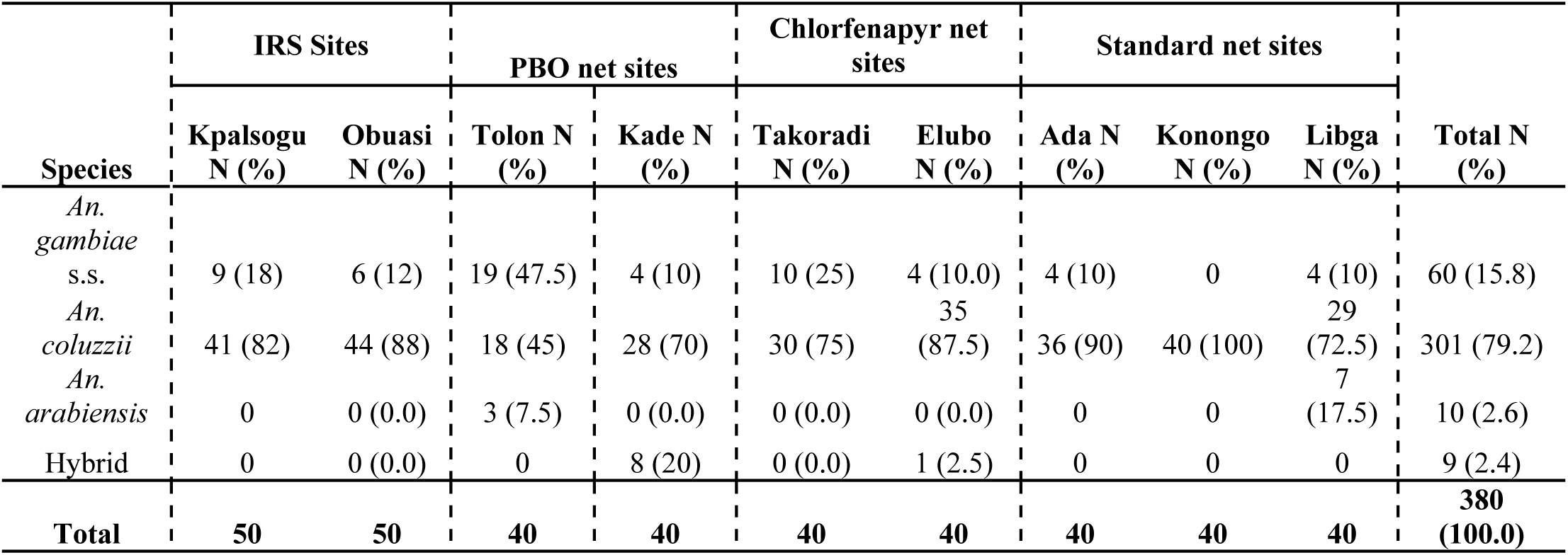
Species discrimination of *Anopheles gambiae* s.l across the study sites.

### Phenotypic Resistance in *Anopheles gambiae* s.l mosquitoes

Moderate to high pyrethroid resistance was observed in all the sites (4% to 65%). Mortalities of *Anopheles gambiae* s.l mosquitoes to permethrin were significantly lower in Obuasi (5%), Dodowa (7.5%) and Takoradi (10%) compared to the remaining sites (χ2 = 146.109, *df* = 8, *P* <0.001). Resistance to deltamethrin was also detected in all the sites: Obuasi (4%), Takoradi (15%), Elubo (20%), Konongo (28.3%), Ada (31.3%), Libga (32.5%), Tolon (47.5%), Kpalsogu (51.3%), Kade (60%) and Dodowa (60%) (χ2 = 107.37, *df* = 8, *P* <0.001). Significantly lower mortality to pirimiphos methyl was observed in Obuasi (1.7%) compared to the other study sites (χ2 = 179.774, *df* = 8, *P* <0.001) (Fig 2). Significant differences in mortalities to all insecticides were observed across interventions and ecological zones.

**Fig 2:**
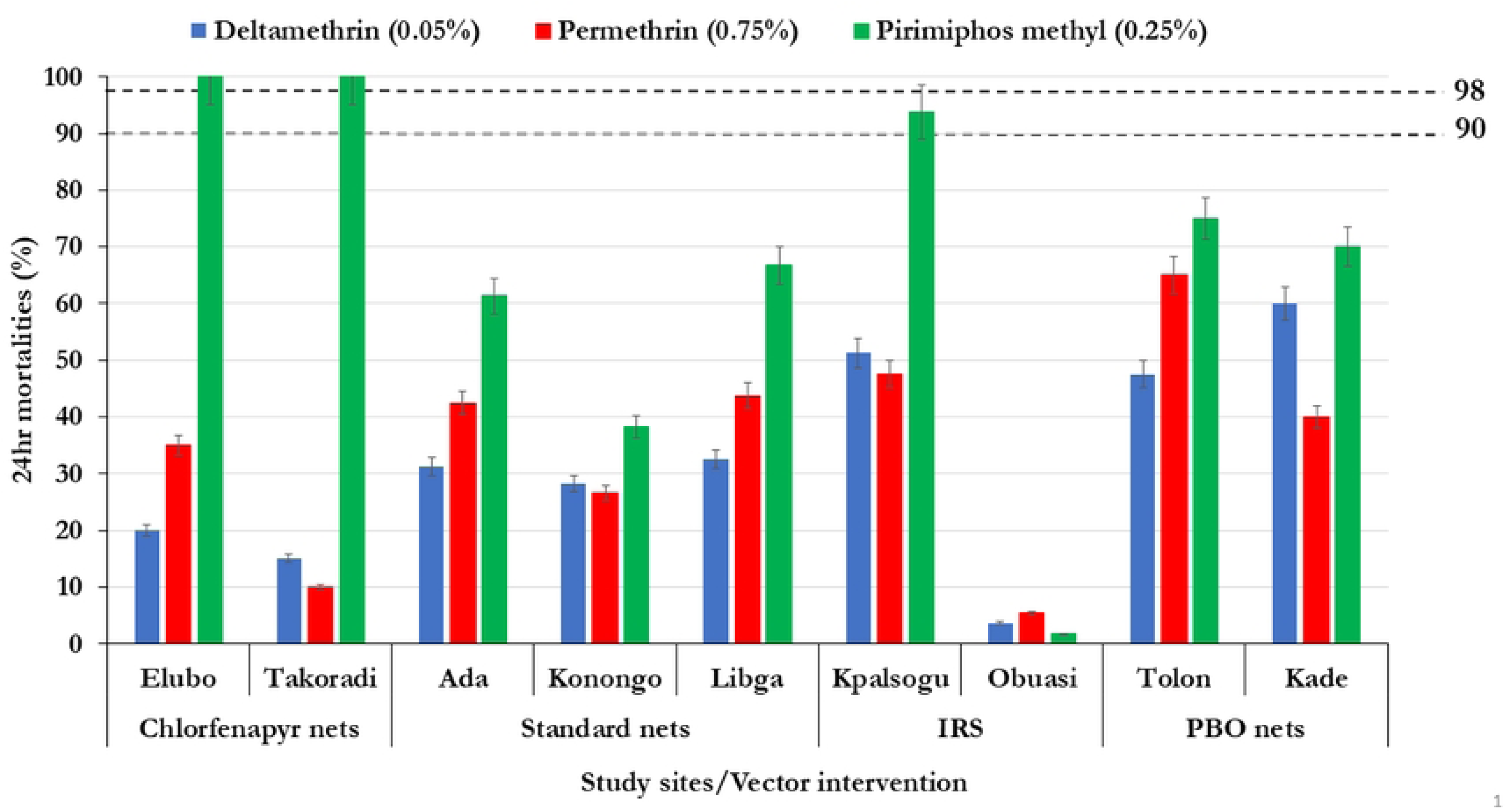
24-hr Mortalities of *An. gambiae* s.l mosquitoes exposed to pyrethroids and organophosphates. Error bars represent the 95% confidence interval of the mean.

### Frequencies of Knockdown Resistance (kdr) Mutations in *An. gambiae* s.l

A subset of 340 pyrethroid phenotyped *An. gambiae* s.l mosquitoes were genotyped for five *kdr* mutations, L995F, L995S, N1570Y, I1527T and V402L. Moderate to High frequencies of the L995F mutation (0.51 -1) was observed in mosquitoes across the study sites, with the highest frequencies found in IRS intervention sites [Obuasi (0.88) and Kpalsogu (0.82)], PBO site Kade (0.88), standard net sites [Konongo (1), Ada (0.89) and Libga (0.83). Lower frequencies of the L995F were observed in *An. gambiae* sl. from PBO net site, Tolon (0.64) and Chlorfenapyr net deployment sites [Elubo (0.51) and Takoradi (0.63)]. Lower frequencies of another kdr mutations, L995S (0-0.03), N1570Y (0 -0.28) and I1527T (0.01 – 0.26) were observed in *An. gambiae* s.l populations from all the study sites. Low to moderate frequencies of the kdr mutation, V402L was observed across the sites. The highest frequency of V402L mutation was observed in *An. gambiae* s.l from Kpalsogu (0.59). Significantly higher frequency of the *kdr* mutation, L995F was observed in pyrethroid resistant mosquitoes (0.85) compared to susceptible ones (0.70) (χ2 = 13.28, *df* = 1, *P*=0.001). The frequencies of other *kdr* mutations between resistant and susceptible mosquitoes were similar with no significant association (P>0.05) (Table 2).

**Table 2:**
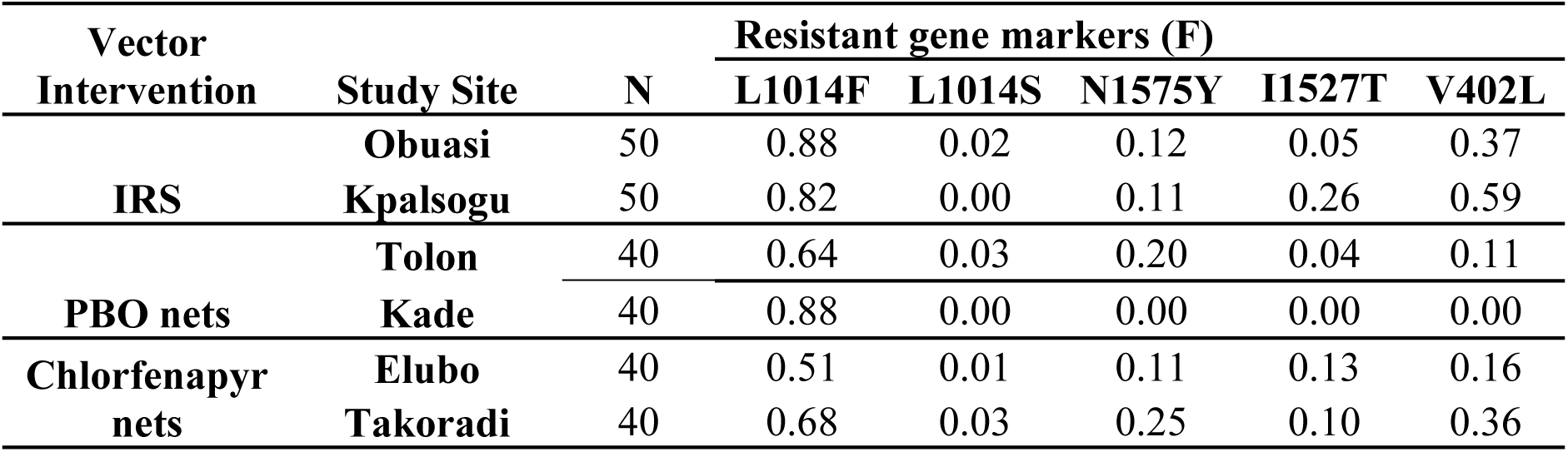
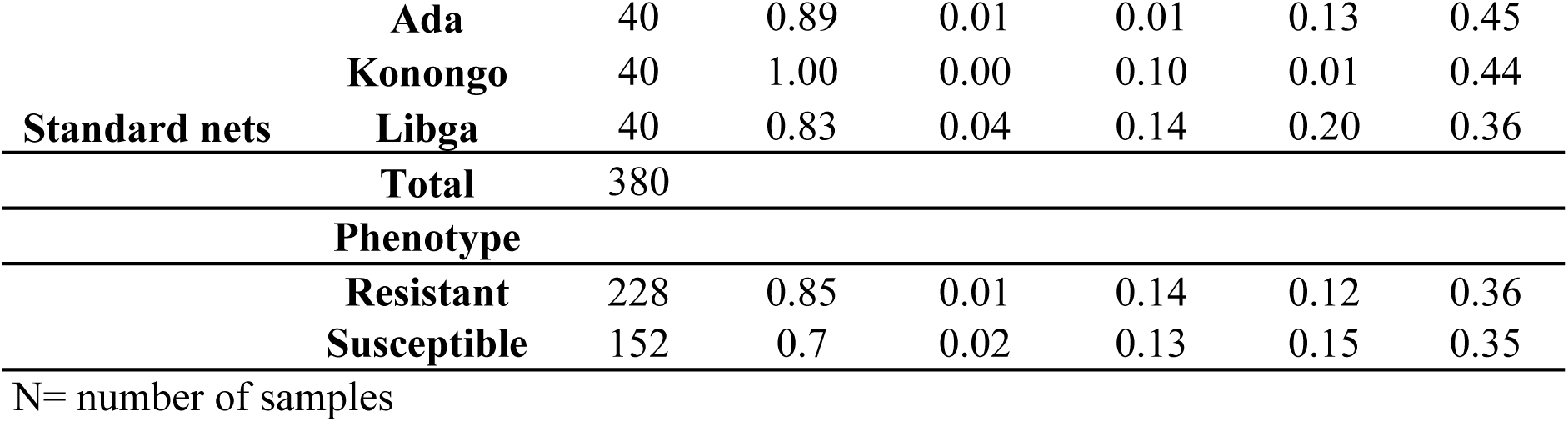
Distribution of knockdown resistance (*kdr*) mutations in *Anopheles gambiae* s.l across the study sites.

### Distribution of *Ace-1* G280S mutation and *Ace-1* gene duplications in *An. gambiae* s.l mosquitoes

A total of 395 *An. gambiae* s.l mosquitoes from six of the study sites phenotyped with organophosphate insecticides were genotyped for the *Ace-1* G280S mutation. The highest frequency of the G280S mutation was observed in *An. gambiae* s.l mosquitoes from standard net sites, Ada (0.44), Konongo (0.38) and Libga (0.36). The least frequency of the G280S mutation was observed in chlorfenapyr net deployment sites, Elubo (0.11) and Takoradi (0.11) (Table 3). There was a significant difference in the frequencies of G280S mutation across the study sites and vector interventions (fishers exact, *P* <0.001). Higher frequency of the *Ace1-* G280S mutation was observed in pirimiphos-methyl resistant *An. gambiae* s.l mosquitoes (0.32) compared to susceptible mosquitoes (0.20) (fishers exact, *P* <0.05). The *Ace1-* G280S mutation was significantly associated with pirimiphos-methyl resistance (OR= 2.20, 95% CI= 1.39 -3.47, *P<0.001*).

**Table 3:** Distribution of *Ace-1* G280S mutation and *Ace-1* gene duplications in *An. gambiae*s.l populations.

| Intervention | Study Site | N | G280S |  |  | Ace-1 | Ace-1 DUP |
| --- | --- | --- | --- | --- | --- | --- | --- |
|  |  |  | RR | RS | SS | F | N (%) |
| IRS | Obuasi | 50 | 2 | 20 | 28 | 0.24 | 6 (12) |
|  | Kpalsogu | 50 | 1 | 20 | 29 | 0.22 | 3 (6) |
| PBO | Tolon | 40 | 4 | 11 | 25 | 0.24 | 11 (27.5) |
|  | Kade | 50 | 0 | 14 | 36 | 0.14 | 1 (2) |
| Chlorfenapyr | Elubo | 40 | 1 | 7 | 32 | 0.11 | 4 (10) |
|  | Takoradi | 40 | 0 | 9 | 31 | 0.20 | 7 (13) |
| Standard | Ada | 50 | 7 | 30 | 13 | 0.44 | 46 (92) |
|  | Konongo | 25 | 6 | 7 | 12 | 0.38 | 5 (20) |
|  | Libga | 50 | 5 | 26 | 19 | 0.36 | 7 (14) |
| Total |  | 395 | 26 | 144 | 225 |  | 90 (22.8) |
| Species |  |  |  |  |  |  |  |
| *An. gambiae* s. s |  | 76 | 2 | 23 | 51 | 0.18 | 8 (10.5) |
| *An. coluzzii* |  | 254 | 16 | 90 | 148 | 0.24 | 70 (27.6) |
| *An. arabiensis* |  | 46 | 6 | 25 | 15 | 0.40 | 7 (15.2) |
| *An. gambiae/coluzzii* hybrids |  | 19 | 2 | 6 | 11 | 0.26 | 5 (26.3) |

F= Allele frequency, N= number of samples, SS = mutant (resistant), GS = heterozygotes, GG= wild type.

A total of 90 (22.8%) *An. gambiae* s.l mosquitoes were found to have Ace-1 gene duplications (*Ace1*_AgDup). The highest frequency of *Ace1*_AgDup was observed in Ada (46/50, 92%) while the lowest frequencies were observed in mosquitoes from Kade (2/50, 2%) (Table 2). There was significant difference in the frequencies of the *Ace1*_AgDup across the study sites and interventions (*P* <0.05). The frequency of the *Ace1*_AgDup was higher in *An. coluzzii* (27.6%) compared to the other species.

A significantly higher percentage of *An. gambiae* s.l mosquitoes with homozygote mutant (61.54%) and heterozygote genotypes (31.94%) had Ace-1 gene duplications compared to homozygote wildtype genotypes (12.44%) (χ2 = 42.73, *df* = 2, *P* =0.000) (Fig 3).

**Fig 3:**
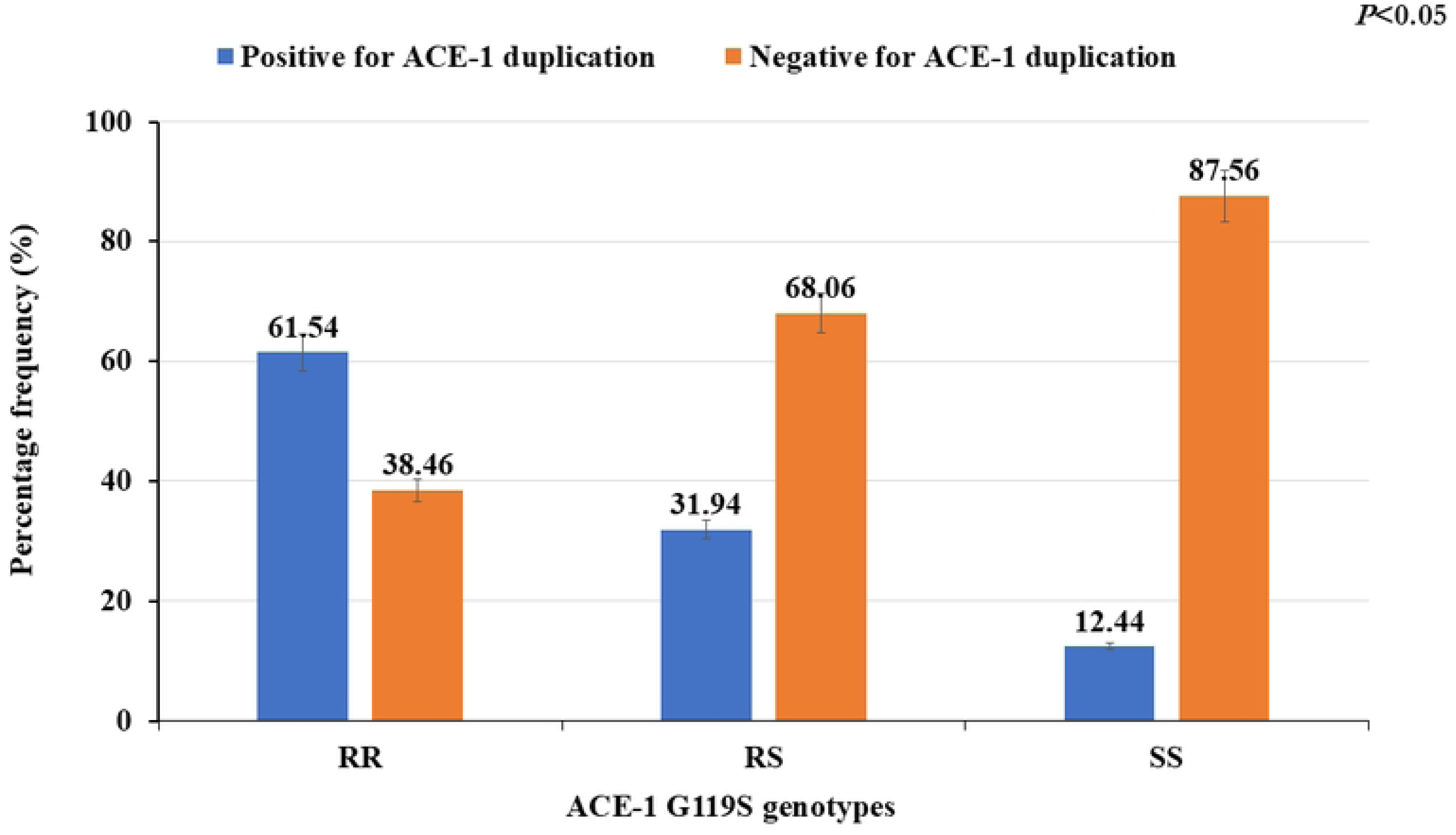
Association between *Ace-1* G280S genotypes and the presence of *Ace-1* gene duplications in *An. gambiae* s.l. Error bars represent the 95% confidence interval of the mean.

### Distribution of Carboxylesterase gene duplications in *An. gambiae s.l*

A total of 395 *Anopheles gambiae* s.l mosquitoes were tested for CNVs spanning the *Coeae1f* and *Coeae2f* genes associated with pirimiphos-methyl resistance. Out of this, 120 (30.4 %) had the duplication junction for Coeaxf_Dup1. The highest frequencies of Coeaxf_Dup1 were found in *An. gambiae* s.l mosquitoes from Ada (80%) and Kpalsogu (52%). A total of 36 (9.1%) *An. gambiae* s.l mosquitoes were positive for Coeaxf_Dup2, with Ada recording the highest frequency (32%) (Table 3). Similar frequencies of Coeaxf_Dup1was found in *An. arabiensis* (39.1%) and *An. gambiae*/*coluzzii* hybrids (42.1%) (Table 4).

**Table 4.** Distribution of *Coeae1f* and *Coeae2f* CNVs in *An. gambiae* s.l across the study sites.

| Intervention | Study Site | N | Coeaxf_Dup1 (%) | Coeaxf_Dup2 (%) |
| --- | --- | --- | --- | --- |
| IRS | Obuasi | 50 | 8 (16) | 2 (4) |
|  | Kpalsogu | 50 | 26 (52) | 1 (2) |
| PBO | Tolon | 40 | 10 (25) | 5 (12.5) |
|  | Kade | 50 | 1 (2) | 3 (6) |
| Chlorfenapyr | Elubo | 40 | 6 (15) | 0 |
|  | Takoradi | 40 | 13 (32.5) | 5 (12.5) |
| Standard | Ada | 50 | 40(80) | 16 (32) |
|  | Konongo | 25 | 2 (8) | 2 (8) |
|  | Libga | 50 | 14 (28) | 2 (4) |
| Total |  | 395 | 120 (30.4) | 36 (9.1) |
| Species |  |  |  |  |
| *An. gambiae* s. s |  | 76 | 10 (13.2) | 3 (3.9) |
| *An. coluzzii* |  | 254 | 84 (33.1) | 30 (11.8) |
| *An. arabiensis* |  | 46 | 18 (39.1) | 1 (2.2) |
| *An. gambiae/ An. coluzzii* hybrids |  | 19 | 8 (42.1) | 2 (9.1) |

N= number of samples

### Associations of CNVs in *Coeae1f/Coeae2f* and *Ace-1* gene with pirimiphos-methyl resistance

We assessed the significance of the association between the presence of CNV copy number of *Coeae1f/Coeae2f* and *Ace-1* genes and their observed phenotype. There was no significant association between the presence of *Ace-1* gene duplication and pirimiphos-methyl resistance despite 0.19 higher odds (OR= 1.19, *P>0.05*). There was a significantly higher odd of resistant *An. gambiae* s.l. having the Coeaxf_Dup1 compared to susceptible mosquitoes (OR= 1.68, *P=0.02*). For the Coeaxf_Dup2, no significant association was observed with resistance phenotypes (*P>0.05*) (Table 5).

**Table 5:**
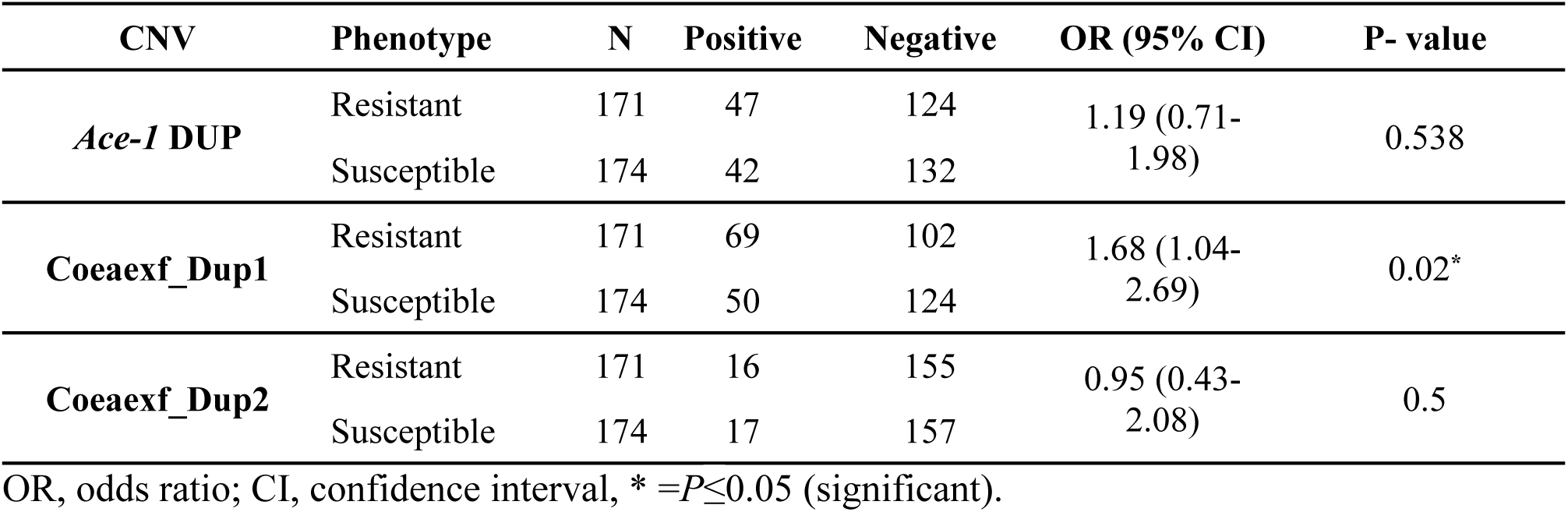
Associations of CNVs with Pirimiphos-methyl Resistance.

## Discussion

Surveillance of insecticide resistance in local malaria vectors is key for effective control and elimination of malaria. This study investigated the distribution of target site resistance mutations and CNVs in *Anopheles gambiae* sensu lato (s.l) mosquitoes from sites with different vector control interventions in Ghana. We detected high pyrethroid resistance in local *An. gambiae* s.l. populations coupled with varying frequencies of insecticide resistance mutations (*kdr* and *Ace-1*). Copy Number Variations (CNVs) in known resistance genes (*Ace-1*, *Coeae1f*, and *Coeae2f*) were also detected in *An. gambiae s.l.* populations. In some cases, these CNVs showed significant associations with insecticide resistance across the study areas.

Our findings revealed high pyrethroid resistance in *An. gambiae* s.l across all the study sites. This may be due to the widespread use of pyrethroids in public health and agriculture pest control (24,38). This widespread deployment of pyrethroid-based control tools such as LLINs have increased the selection pressure for insecticide resistance in malaria vectors. Previous studies from Ghana and other African countries have also documented evidence of pyrethroid resistance in *Anopheles gambiae* s.l mosquitoes (24,28,39–41).

Resistance to the organophosphate, pirimiphos-methyl was observed in *An. gambiae* s.l from all sites with the exception of Chlorfenapyr net deployment sites, Elubo and Takoradi. This shows that pirimiphos-methyl resistance is increasing in local *An. gambiae* s.l populations. This may be due to the scale-up in the use of pirimiphos methyl for indoor residual spraying by the National Malaria Elimination Program (NMEP) in some sites in Ghana to control wide spread pyrethroid resistance (38). This may explain why *An. gambiae* s.l. mosquitoes from the IRS site, Obuasi had very high resistance to pirimiphos-methyl. Moreover, the use of organophosphates for agricultural pesticides may also contribute to the high resistance observed in mosquitoes from the non-IRS sites such as Ada and Konongo (24,39). Other studies in Ghana have also detected resistance to pirimiphos-methyl in malaria vectors(23,24). This highlights the need for increased monitoring of organophosphate and carbamate resistance in *An. gambiae* s.l populations in Ghana.

Moderate to high frequencies of the *kdr* mutation, L995F was observed across the sites. *Anopheles gambiae* s.l from IRS and standard net deployment sites had significantly higher frequencies of L995F mutation compared to sites where newer next generation nets are deployed. High frequencies of L995F mutation have also been observed in other studies across Ghana(12,42,43). Lower frequencies of the L995S, N1570Y and I1527T were observed in *An. gambiae* s.l from all the sites, similar to findings from other studies (12,16,44). Previous studies reported a significantly lower frequencies of these I1527T and V402L in Ghanaian Anopheles populations compared to findings from this study(16). Findings from this study showed low to moderate frequencies of V402L (0.11 to 0.59), similar to findings from other studies(17,45).

The *Ace-1* G280S mutation was significantly associated with pirimiphos methyl resistance, similar to findings from other studies across West Africa(17,23). The *Ace-1* G280S was found at low to moderate frequencies, similar to frequencies observed in local *An. gambiae* s.l populations by Hamid Adiamoh *et al*.(12) and Owusu-Asenso et al.(46). In Contrast, a study in Accra, Southern Ghana reported very high frequencies of the *Ace-1* G80S mutation (0.76) in *An. gambiae* s.l mosquitoes(24). A total of 22.8% of *An. gambiae* s.l mosquitoes were found to have two or more copies of the *Ace-1* loci, which is higher than a previous study by Grau-Bové *et al*.(23), where frequencies of 0.88 to 0.11 was detected in *An. gambiae* s.l populations from Ghana. Previous studies have reported associations between the *Ace-1* CNV and resistance to organophosphates(37,47). In contrast, our results did not reveal a significant association to pirimiphos-methyl resistance. This may be due to the limited sample size used in the study.

Over the years, organophosphate resistance was widely attributed to the *Ace-1* locus, where gene duplications coupled with the *Ace-1* G280S mutation contributes to variable levels of resistance (23,37). However, metabolic and cuticular mechanisms have also been implicated in organophosphate resistance(48,49). Metabolic genes *Coeae1f*, and to a lesser extent *Coeae2f*, are frequently overexpressed in insecticide-resistant populations compared to susceptible ones in sub-Saharan Africa(21). Our findings revealed the presence of CNVs of metabolic gene locus, *Coeae1f/2f* in local *An. gambiae* s.l mosquitoes. We found *Coeae1f* CNVs in about 30.4% of the study population. The first report of the *Coeae1f/2f* CNVs was by Nagi *et al*.(21), where they detected CNVs in *Coeae1f* at frequencies of 14% in Obuasi, similar to the 16% rate detected in *An. gambiae* s.l from Obuasi in this study. A significantly high frequency of the *Coeae1f* was found in Ada (80%) compared to the other sites. Ada is an irrigated farming site where standard pyrethroid nets are deployed. It is possible that other selective pressures from agricultural pesticides may be contributing to the high frequencies of CNVs associated with organophosphate resistance observed from our study.

Our findings also revealed that the CNVs in *Coeae1f* was significantly associated with pirimiphos-methyl resistance, in line with study by Nagi *et al*.(21). Although the observed association is consistent with previous findings, its interpretation is limited by the assay’s detection of a duplication junction rather than direct evidence of the CNV’s functional effect. While these findings suggest that Coeae1f may represent a promising candidate locus associated with pirimiphos-methyl resistance in *An. gambiae* s.l. from Ghana, they do not independently establish a causal role or resistance mechanism. Additional studies, including functional validation, are needed to confirm its contribution to insecticide resistance. Pirimiphos-methyl, formulated as Actellic CS300, was widely used for several years in IRS campaigns in Ghana(38) and throughout sub-Saharan Africa(50,51). Hence it is important to increase surveillance of newly emerged resistance mechanism to help in insecticide resistance management.

The predominant species across the study sites was *Anopheles coluzzii*, similar to finding from other studies in Ghana(42,52,53). *Anopheles coluzzii* is commonly breeds in permanent breeding sites, hence its dominance in most of the study sites(52,54). *Anopheles arabiensis* was only found in the Sahel sites, Libga and Tolon, in line with previous studies in Ghana, where it has been found exclusively in the Sahel zone because of its adaptation to arid environments(52,53,55).

## Conclusion

This study revealed widespread insecticide resistance to pyrethroids across all the sites. Frequencies of *kdr* and *Ace-1* gene mutations varied across the sites. Copy Number Variations (CNVs) of known insecticide resistance genes (*Ace-1*, *Coeae1f* and *Coeae2f*) was also detected in *An. gambiae* s.l. populations, with some showing significant associations with insecticide resistance across the study sites. This highlights the need for more surveillance and research into current and emerging insecticide resistance mechanisms to help improve current malaria vector control efforts in Ghana.

## Acknowledgement

We thank all the residents within the study sites for their support during the study. Our since gratitude to the entire team of the Centre for Vector-borne Disease Research (CVBDR) team at the Department of Medical Microbiology, University of Ghana Medical School, for assisting with the field and laboratory work.

## Notes

### Competing Interest Statement

The authors have declared no competing interest.

